# Functional analysis of FW2.2-LIKE proteins in tomato: SlFWL5 regulates leaf size and morphology

**DOI:** 10.1101/2025.02.27.640525

**Authors:** Arthur Beauchet, Lucie Ehrhard, Lina Boutaleb, Valérie Rofidal, Nathalie Gonzalez, Christian Chevalier, Norbert Bollier

## Abstract

The *CELL NUMBER REGULATOR/FW2.2-like* (*CNR/FWL*) gene family comprises PLAC8 domain-containing membrane-associated proteins. The *CNR/FWL* gene family was named in reference to its founding member, the *FW2.2* gene, which determines fruit size in tomato via a negative regulation on cell divisions. The functional characterization of PLAC8 domain-containing CNR/FWL proteins in plants is still very scarce, and only recently the molecular and cellular mode of action of FW2.2 was identified as regulating plasmodesmata-mediated cell-to-cell communication. In the present study, we provided a functional analysis of *SlFWL* genes in tomato, aiming at investigating any direct role in the control of organ growth. Based on a combination of molecular and cellular approaches, we selected three SlFWL proteins, namely SlFWL2, -4 and -5, which are localized at the plasma membrane. Gain- and loss-of-function transgenic plants were generated to explore their putative role as regulators of organ growth in tomato. This allowed us to shed light more specifically on the critical involvement of SlFWL5 in leaf development. We here showed that SlFWL5 is a plasmodesmata-localized protein that regulates leaf size and morphology via cell expansion, most probably through cell-to-cell communication. This original finding underscores further the importance of the CNR/FWL family members in growth regulation.

**Highlight:** The FW2.2-LIKE protein SlFWL5 is localized at plasmodesmata and may have a role in cell-to-cell communication to regulate cell expansion during leaf development.

## Introduction

The *CELL NUMBER REGULATOR/FW2.2-like* (*CNR/FWL*) gene family (Guo *et al*., 2010; Thibivilliers *et al*., 2020) was named in reference to its founding member, the *FW2.2* gene (standing for Fruit Weight QTL on chromosome 2, number 2), which underlies the major quantitative trait locus (QTL) governing fruit size in tomato (Alpert *et al*., 1995; Frary *et al*., 2000). The *CNR/FWL* gene family encompasses hundreds of related sequences in the plant, animal, and fungal reigns (Guo *et al*., 2010), with a large diversity in protein size ranging from a hundred to several hundred amino acids. The CNR/FWL proteins are usually described as membrane-associated Pro- and Cys rich proteins, which harbor as a principal feature, the presence of a conserved structural domain named PLAC8 (Placenta specific 8), originally identified in a protein specifically expressed in the trophoblast giant cells and spongiotrophoblast layer of the placenta from mouse (Galaviz-Hernandez *et al*., 2003). The PLAC8 domain is composed of one or two hydrophobic segments, predicted to form two putative transmembrane (TM) α-helices, and characterized by the presence of Cys-rich motifs of the CLXXXXCPC or CCXXXXCPC type (Song *et al*., 2004; Beauchet *et al*., 2021).

Three main functions have been assigned to PLAC8 domain-containing CNR/FWL proteins in plants: the regulation of organ size, especially in fruits, cadmium resistance, and metal ion homeostasis (Beauchet *et al*., 2021). Indeed, the original FW2.2 protein regulates negatively cell divisions in the pre-anthesis ovary and young developing fruit, thereby accounting for its quantitative effects on final fruit size via the modulation of cell number (Nesbitt and Tanksley, 2001; Cong *et al*., 2002; Liu *et al*., 2003). Functional analyses of CNR/FWL proteins suggest that this role in the regulation of organ growth, primarily through their effects on cell division, is a highly conserved feature in both monocotyledon and dicotyledon plants (Dahan *et al*., 2010; Guo *et al*., 2010; Li and He, 2015; Qiao *et al*., 2017; Ruan *et al*., 2020). Interestingly and related to this function, the PLAC8 protein has been determined as well to be involved in organ development and cell division control during tumorigenesis in mammals (Cabreira-Cagliari *et al*., 2018; Mao *et al*., 2021). However, the precise molecular mechanism underlying the role of CNR/FWL proteins in regulating organ growth linked to cell division/cell cycle control remain largely unexplored.

Recently, Beauchet *et al*. (2024) made a major breakthrough towards the identification of the molecular and cellular mode of action of FW2.2 in tomato. In this study, FW2.2 was demonstrated to be a membrane-anchored protein that locates at plasmodesmata (PD) and regulates cell-to-cell communication by modulating PD transport capacity during fruit development, via the regulation of callose deposition at PD (Beauchet *et al*., 2024). Whether this function is conserved in CNR/FWL proteins from other plant species remains to be determined. Nevertheless, several clues seem to comfort this functionality. Indeed, the closest ortholog of FW2.2 in Arabidopsis, namely AtPCR2, is highly enriched at PD (Brault *et al*., 2019). The soybean GmFWL1 and GmFWL3 proteins are localized to membrane microdomains, and interacts with structural proteins of the membrane microdomains, but also with proteins involved in the metabolic process of callose deposition, such as callose synthases (Qiao *et al*., 2017; Cervantes-Pérez *et al*., 2024), alike FW2.2 in tomato fruit (Beauchet *et al*., 2024).

An updated phylogenetic analysis performed on 13 different plant genomes released in recent years allowed the identification of 134 members in the CNR/FWL family, with a total number of twelve *FWL* genes including *FW2.2* in tomato (Thibivilliers *et al*., 2020). This list has been recently extended up to 21 members (Ran *et al*., 2023), providing a valuable resource for the characterization of *FWLs* in tomato.

In the present study, we provided the first functional analysis of *SlFWL* genes in tomato. We aimed at investigating whether SlFWLs can play a direct role in the control of organ growth. First, we re-investigated the expression profiles of *SlFWLs* in different organs of tomato plants and during fruit development, and analyzed their subcellular localization. Based on this combination of molecular and cellular approaches, we selected three *SlFWL* genes, namely *SlFWL2*, -*4* and -*5*, and generated gain- and loss-of-function transgenic plants to explore their putative role as regulators of organ growth and development in tomato. This allowed us to shed light more specifically on the critical involvement of SlFWL5 in the regulation of leaf development via cell expansion, further underscoring the importance of the CNR/FWL family members in growth regulation.

## Materials and methods

### Plant materials and growth conditions

Tomato (*Solanum lycopersicum* cv. AC) and tobacco (*Nicotiana benthamiana*) plants were grown in soil in a greenhouse under the following conditions: 16 h day/8 h night cycle, using a set of 100 W warm white LED projectors providing an irradiance of 100 μmol m^−2^ s^−1^ at the level of canopy. The light spectrum was constituted by equivalent levels of blue irradiation (range 430–450 nm) and red irradiation (640–660nm). For in *vitro* culture, tomato seeds were sterilized for 10 min under agitation in a solution of 3.2% bleach. Seeds were then washed three times with sterile water and dried under a laminar flow hood. Seeds were sowed in Murashige and Skoog medium (1/4 MS) and transferred in a growth chamber under the following conditions: 16 h day/8 h night cycle, 22°C/20°C day/night, using white light (Osram L36 W/77 Fluora 1400 Im) providing 80 to 100 μE m^−2^ s^−1^ intensity light at the stirring plate.

### Phylogenetic analysis and tools for the prediction of the CNR/FWL structure and topology

The 21 SlFWL protein sequences from tomato were mined from Thibivilliers *et al*. (2020) and Ran *et al*. (2023), and aligned using MEGA11 Muscle alignment (Tamura *et al*., 2021). Values for 1000 bootstrap replicates are shown on each branch.

DeepTMHMM (https://dtu.biolib.com/DeepTMHMM) (Hallgren *et al*., 2022) was used to predict the presence of transmembrane helix in SlFWL proteins. The 3-dimensional structure of the full-length SlFWL2, -3 and -5 proteins was obtained using Colabfold (Mirdita *et al*., 2022) The PPM 3.0 Web Server (https://opm.phar.umich.edu/ppm_server3_cgopm) (Lomize *et al*., 2022) was used with default parameters and plasma membrane (plants) type to predict the topology and insertion of SlFWL2, -3 and -5 in the plasma membrane.

### Vector constructs and plant transformation

Vectors for the overexpression of *SlFWLs* in plants were generated using the Gateway® cloning system (Invitrogen, Carlsbad, CA, USA), following manufacturer’s instruction. The *FWLs* full-length coding sequence was amplified from cDNAs prepared from various tomato tissues (cv. AC) using PrimeSTAR MAX DNA polymerase (TAKARA BIO Inc., Kusatsu, Japan) and primers including the attB sites (**Supplemental Table 1**). The resulting PCR products were cloned into the corresponding Gateway vectors described in **Supplemental Table 2**. For CRISPR/Cas9 mutagenesis, constructs were assembled using the Golden Gate cloning method according to previous method (Beauchet *et al*., 2024). Transgenic plants were generated by *Agrobacterium tumefaciens* (strain C58C1) mediated transformation using explants of tomato cotyledons as described (Swinnen *et al.,* 2022).

### RNA extraction and RT-qPCR analysis

Total RNA was isolated from cotyledons, hypocotyls, shoot apical meristems, leaves, roots, flowers and pericarp tissues from fruits harvested at different developmental stages (5, 10, and 15 DPA), using TRIzol reagent (Invitrogen) in combination with RNeasy Plant Mini Kit (Qiagen) following the manufacturers’ instructions. RNase-free DNase (Qiagen) treatment was performed on each sample. Reverse transcription was performed using the iScript^TM^ cDNA Synthesis Kit (Bio-Rad, Hercules, CA). Real-time PCR was performed using Gotaq® qPCR mastermix (Promega, Madison, WI) and a CFX 96 real-time system (Bio-Rad). qPCR primers were designed with PerlPrimer software (Marshall, 2004) to overlap 2 exons in order to limit genomic DNA amplification (**Supplemental Table 2**) and amplify a 80 to 200 bp-long amplicon, with a Tm of 60°C. The transcript levels of the expressed genes were normalized to that of the housekeeping genes: *SlTUBULIN* (Solyc04g081490) in combination with *SlNUDK* (Solyc01g089970) for fruit samples, or with *SlEIF4α* (Solyc12g095990) for other tissue samples.

### Subcellular localization analysis of SlFWL proteins

The subcellular localization of SlFWLs fused to YFP was observed using confocal imaging performed on a Zeiss LSM 880 confocal laser scanning microscope equipped with fast AiryScan, using a Zeiss Plan apochromat 10x/0.45 or a C PL APO x63/1.4 oil-immersion objective.

The top three leaves of three independent *N. benthamiana* plants (6–8 weeks old) were agro-infiltrated with *35S::SlFWLs-YFP* constructs and plasmolyzed using 0.4 M Sorbitol for 15 min before observation. Co-infiltrations with SP-mTagBFP2-HDEL as a marker of ER and mScarlet-l as a marker of both cytosol and nucleus localization were performed to ascertain the localization pattern. Staining with FM4.64 at a final concentration of 4 µM was used as a control for PM localization (Bolte *et al*., 2004). For FM4.64 imaging, excitation was performed at 561 nm and fluorescence emission was collected at 630-690 nm. For YFP imaging, excitation was performed at 514 nm and fluorescence emission collected at 520-580 nm. For mTagBFP2 imaging, excitation was performed at 405 nm and fluorescence emission was collected at 440-490 nm. For mScarlet-I imaging, excitation was performed at 561 nm and fluorescence emission was collected at 570-640 nm.

The localization of SlFWL2, -4 and 5 at PM and PD was performed in leaves of tomato *35S::SlFWL2-YFP*, *35S::SlFWL4-YFP* and *35S::SlFWL5-YFP* transgenic plants. Similarly, leaf epidermal cells were plasmolyzed using 0.4 M Sorbitol prior to observation. Staining with aniline blue (Biosupplies, Victoria, Australia) was performed by infiltration of a 0.0125% solution; excitation was performed at 405 nm and fluorescence emission collected at 420-480 nm. The calculation of PD index was determined by calculating the fluorescence intensity of SlFWL2-YFP, SlFWL4-YFP and SlFWL5-YFP at plasmodesmata and at PM as described previously (Grison *et al*., 2019).

### Phenotypic characterization

Plants were cultivated randomly side-by-side with WT plants. Flowers were vibrated every day to ensure optimal self-pollination. Seven flowers per inflorescence were maintained to ensure proper development of fruit per inflorescence. Fruits from four to six plants of each genotype of two biological replicates were used to determine fruit weight at the breaker stage of fruit development. Fruits were weighted and measured using a caliper. The number of measurements ranged from n= 50 to n= 200 depending on the number of fruits produced by the different transgenic plants. For root and hypocotyl length measurements, seedlings were grown on MS1/4 in 245 x 245 x 20 mm Nunc tissue culture plates placed vertically. Pictures of seedlings were digitized using a scanner every day and root and hypocotyl length were measured using ImageJ software. For leaf surface phenotyping, pictures of full-grown leaves were taken using a Nikon D5300 and analysed by intensity threshold filtering in ImageJ. For leaf epidermal cell size analysis leaflets were cleared with 70% ethanol for several days before staining with Calcofluor white M2R stain 0.1 % (w/v) for 48h. Leaflets fragments were observed with a Zeiss Axioimager epifluorescence microscope equipped with a 20x (NA 0.5) using a DAPI-BP filter and a Cool Snap HQ2 CCD Camera.

### Co-immunoprecipitation and mass-spectrometry analysis

Total protein extracts from 100 mg of *35S::SlFWL5-YFP* leaf tissue were prepared using the following buffer: 1X PBS, cOmplete Protease Inhibitor Cocktail tablets (Roche, Mannheim, Germany) and 1% Triton X-100. Samples were incubated in the extraction buffer at 4°C for 30 min with agitation, and then centrifuged (16000*g*, 10 min, 4°C). Prior to co-immunoprecipitation, western-blotting was used to check the presence of the expressed tagged-SlFWL5-YFP protein in the supernatant. The supernatant containing the resuspended proteins was then used for immunoprecipitation assay using anti-GFP microbeads provided in the μMACS Epitope Tag Protein Isolation Kit according to the manufacturer’s protocol (Miltenyi Biotec, Bergisch Gladbach, Germany). Protein concentration was determined by Bradford assay according to the manufacturer’s instructions (Pierce Coomassie Plus (Bradford) Assay kit; Thermo Scientific). Approximately, 500 μg of soluble proteins were loaded for each co-IP assay.The LC-MS-MS experiment was carried out as reported previously (Beauchet *et al*., 2024).

The GO enrichment analysis was carried out on PLAZA 5.0 (Van Bel *et al*., 2022) using the Plaza workbench, with a significance threshold of 0.05 and without any data filter.

## Results

### Phylogenetic classification and gene expression profiles of SlFWLs

A recent genome-wide analysis of the *SlFWL* family identified a total of 20 *FW2.2* homologous sequences in tomato (Ran *et al*., 2023). These authors named the different genes according to their location on the tomato chromosomes, not taking into account the anteriority of the *SlFWL* gene classification (although incomplete) proposed by Thibivilliers *et al*. (2020). To avoid additional confusion and to compel to the principle of anteriority, we proposed to keep the latter gene annotation and to assign ascending gene names to the 9 new identified SlFWL sequences according to their phylogenetic relationship to FW2.2 (**Table I**).

**Table I.**
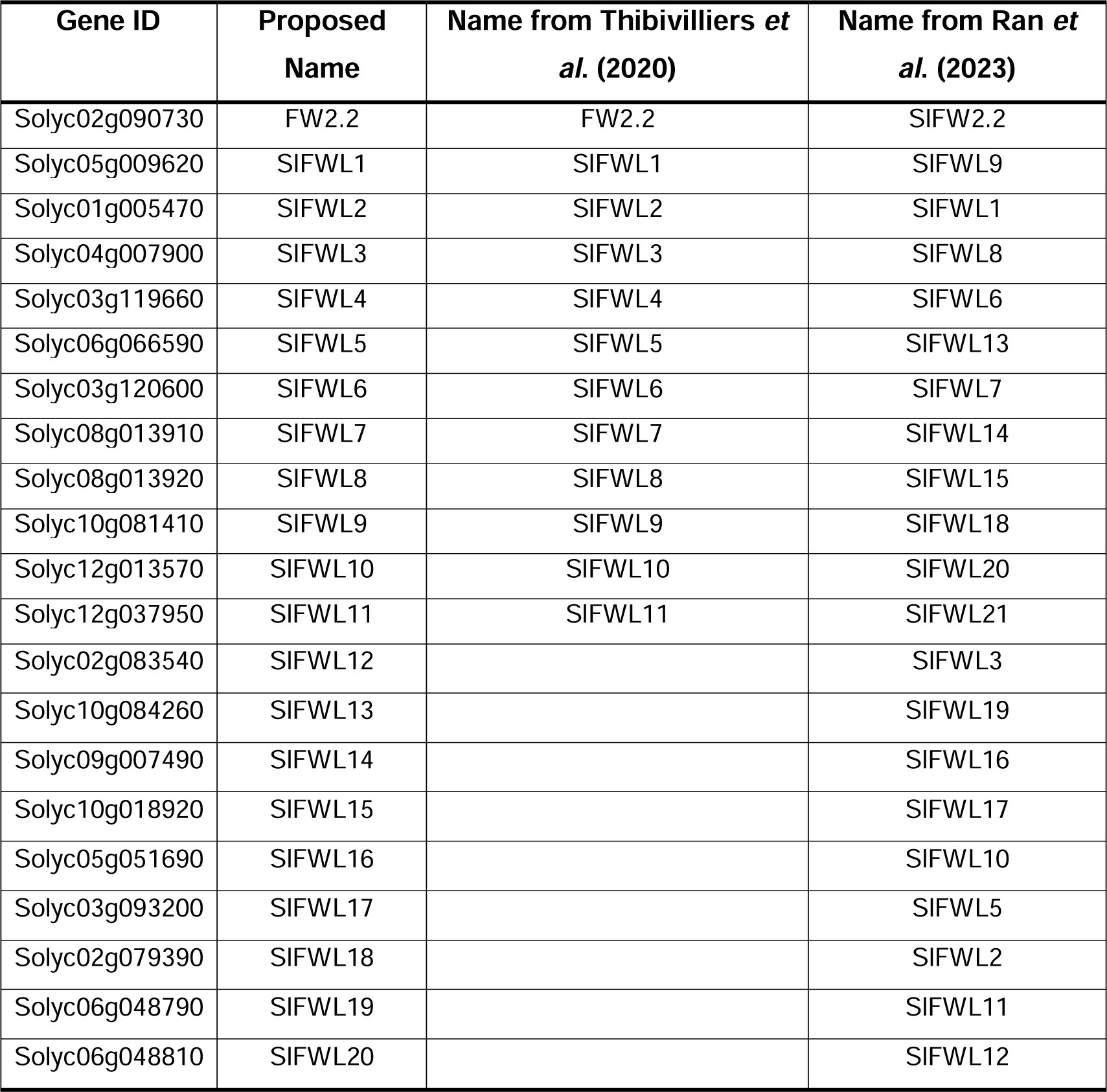
Proposed nomenclature of *Solanum lycopersicum FWL* genes.

A phylogenic tree was thus generated using the full-length sequences of FW2.2 and the 20 homologous sequences from tomato (**Figure 1A**). The SlFWLs fall into two distant clades: the first clade includes FW2.2 and the eleven sequences identified by Thibivilliers *et al*. (2020), with the addition of SlFWL12; the second clade encompasses SlFWL13 to SlFWL20. Remarkably, SlFWLs from the first clade display a highly similar protein homology, with the exception of SlFWL4, SlFWL7, -8 and -12 which harbor a long N-terminal extension upstream of the conserved PLAC8 domain. SlFWLs from the second clade differ from the first clade by a longer sequence, a greater distance between the two conserved motifs of the PLAC8 domain and/or the presence of two or more predicted transmembrane (TM) α-helices as revealed by the use of the transmembrane topology prediction tool DeepTMHMM (Hallgren *et al*. 2022) (**Figure 1A**). It is noteworthy that SlFWL9 occupies an intermediate position between the two subgroups, with a relatively small sequence than that of FW2.2 and the presence of two TM domains.

**Fig. 1.**
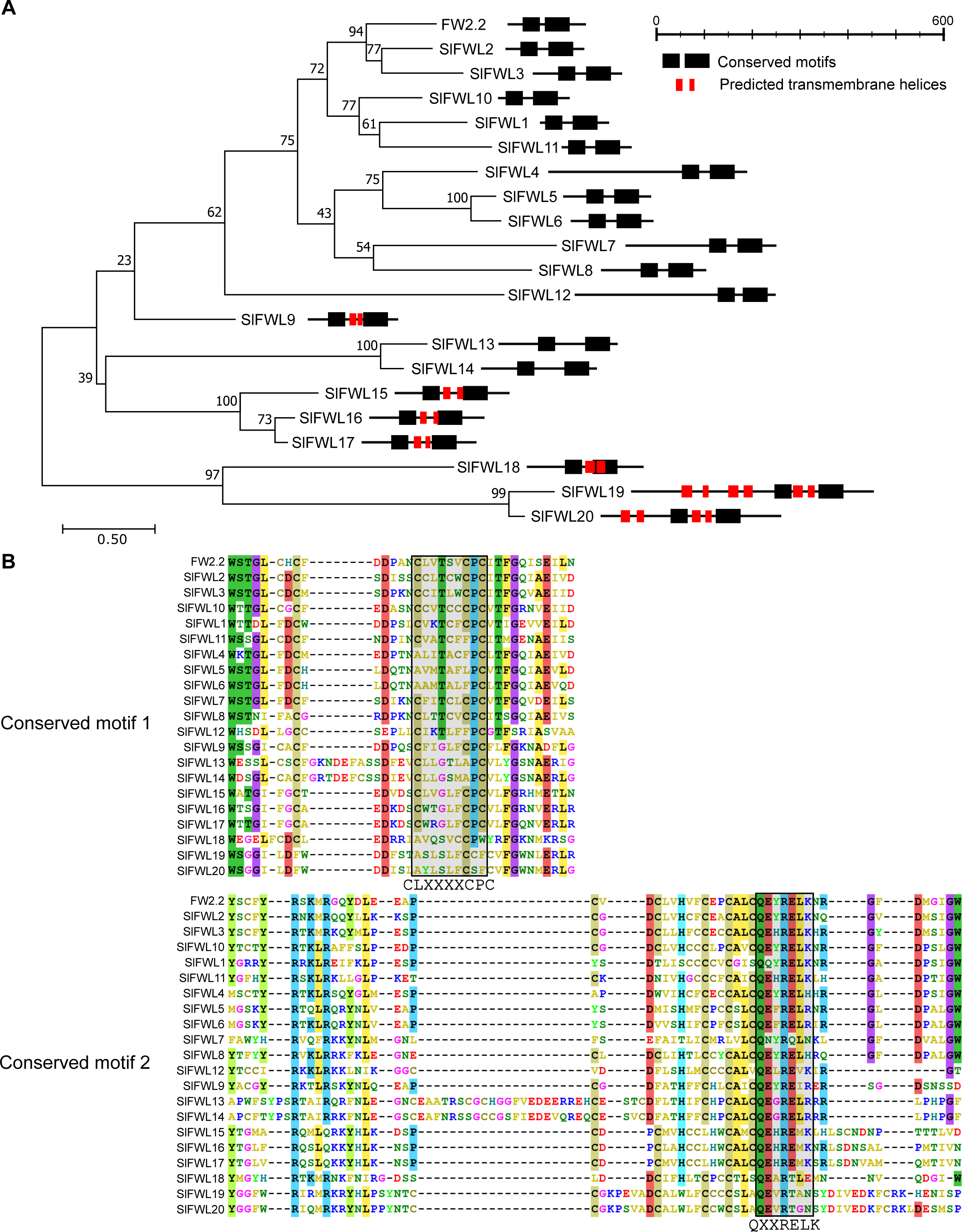
Phylogenetic analysis and protein structure of SlFWLs. (A) Phylogenetic tree built using maximum likelihood method was generated using MEGA11 (Tamura *et al*., 2021). Values for 1000 bootstrap replicates are shown on each branch. The schematic representation of the SlFWLs protein structure were deduced from computation analysis of protein primary sequences. Transmembrane domains were predicted using DeepTMHMM (https://dtu.biolib.com/DeepTMHMM) (Hallgren *et al*., 2022). (B) Amino acid sequence comparison of the PLAC8 domain between SlFWLs and FW2.2, using Muscle alignment (MEGA11).

**Figure 1B** describes the primary structure of the conserved PLAC8 domains in the 20 SlFWL proteins. The CLXXXXCPC motif is only conserved in FW2.2, SlFWL8 and SlFWL15, while SlFWL13 and -14 harbors a CLXXXXAPC sequence. SlFWL2, -3 and -10 harbors the CCXXXXCPC sequence. The CLXXXXCPC or CCXXXXCPC motif diverges more or less in the other SlFWL proteins, especially for the two first amino acids of the sequence. The second motif at the C-terminal part, the so-called ‘QXXRELK’ motif, appears much more conserved among the 20 SlFWL proteins.

To provide an overview of the expression pattern in tomato, the levels of transcript accumulation for *SlFWL* genes were monitored by RT-qPCR in various organs and fruit developmental stages from the tomato cultivar ‘Ailsa Craig’ (‘AC’) (**Supplemental Figure S1**). This analysis showed that a large diversity of expression profiles was observed among the tested *SlFWL* genes, not only in terms of organs, but also in terms of level of transcript accumulation. For instance, *SlFWL3*, *SlFWL14*, and *SlFWL15* exhibited much higher expression levels, ∼a hundred-fold more, than all the other *SlFWL* genes. Most of the *SlFWL* genes are expressed to various levels throughout fruit development. *SlFWL3*, *SlFWL9*, *SlFWL910* and *SlFWL15* were predominantly expressed in fruits, with *SlFWL3* and *SlFWL15* having the highest expression. *FW2.2*, *SlFWL1* and *SlFWL4* were highly expressed in roots and to a lesser extent in fruit. *SlFWL2* was predominantly expressed in cotyledons, and to a lesser extent in hypocotyls, leaves and flowers, and slightly in roots. *SlFWL5* showed high expression in both roots, cotyledons and to a lower extent in leaves and hypocotyl. *SlFWL7* and *SlFWL8* were more specifically expressed in flowers. *SlFWL13*, *SlFWL14*, *SlFWL16*, *SlFWL17* and *SlFWL18*, were broadly expressed across nearly all tissues.

### Sub-cellular localization of SlFWL proteins

The subcellular localization of SlFWL proteins was first investigated in *Nicotiana benthamiana* leaves, using transient expression assays. Constructs aimed at expressing the SlFWL proteins fused to YFP at the C-terminal end, under the control of the Cauliflower Mosaic Virus (CaMV) 35S promoter were thus generated (referred to as *35S::SlFWLX-YFP*). This type of constructs was chosen based on our previous work, showing that the sub-cellular localization of FW2.2 was independent from the position of YFP at the C-terminal or N-terminal end of the protein (Beauchet *et al*., 2024). **Figure 2** displays the data obtained for eleven out of twenty SlFWLs. The following SlFWLs were not included in this analysis for the following reasons: (i) we could not state in the subcellular localization of SlFWL3, due to a lack of reproducibility in the results; (ii) we were unable to generate the constructs fusing the coding sequence for SlFWL12, -13, -14, -16, -17 to YFP for cloning technical reasons; (iii) SlFWL18, -19 and -20 were excluded from the study, because of their too distant phylogenetic relationship with the other SlFWLs.

**Fig. 2.**
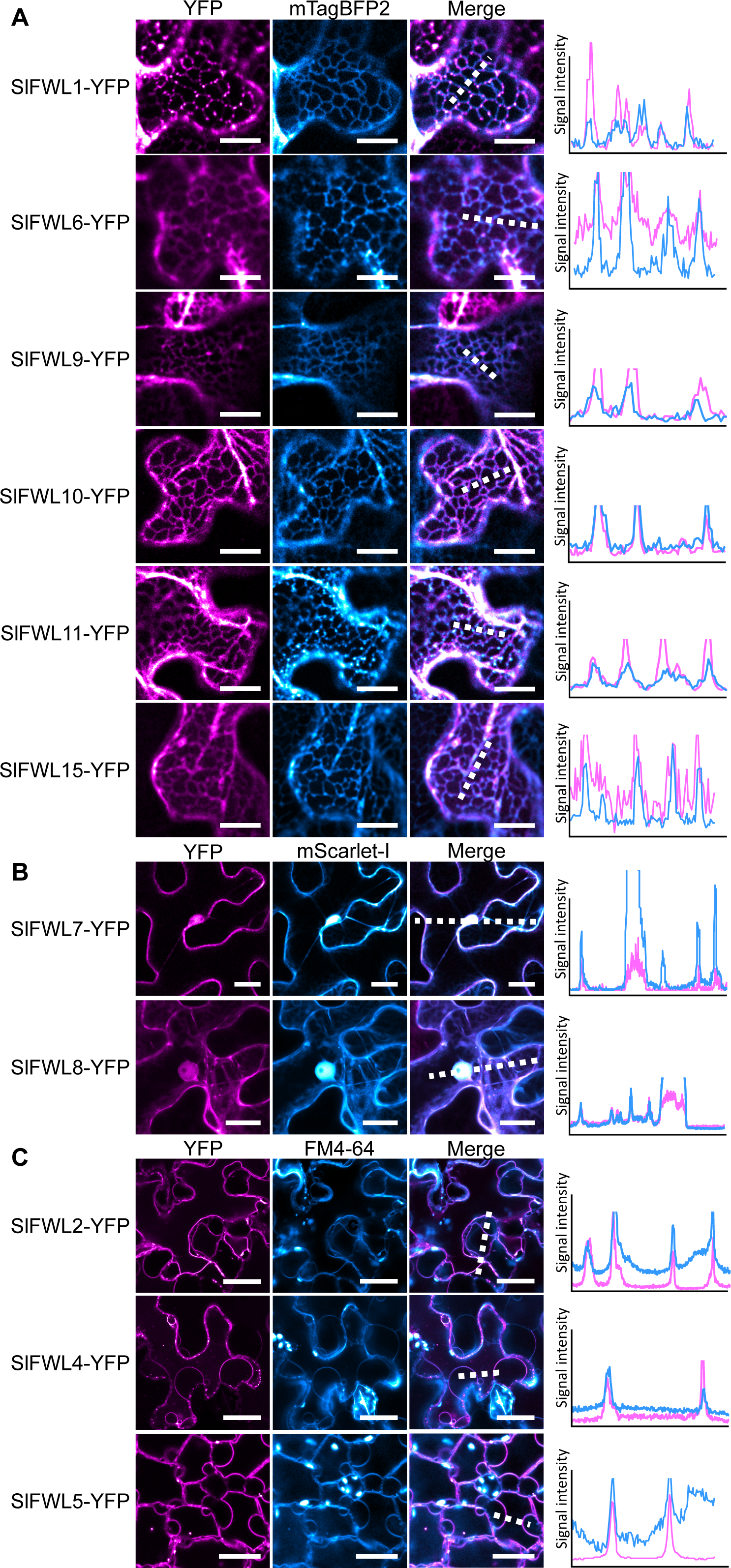
Subcellular localization of SlFWLs fused to YFP in N. benthamiana leaf epidermal cells. (A) ER localization witnessed by the use of the ER-specific marker construct *pUBI::mTagBFP2-HDEL*. Scale bar = 10µm. (B) Cytosol and nucleus localization witnessed by the use of the marker construct *pUBI::mScarlet-I*. Scale bar = 10µm. (C) PM localization witnessed by the membrane specific dye FM4-64. Scale bar = 20µm. Intensity plots delineated by the dashed lines are shown for each co-localization pattern.

For the tested SlFWLs, three distinct patterns of subcellular localization were observed. First, SlFWL1, -6, -9, -10, -11 and -15 were distributed within the cell in a reticular pattern, consistent with a localization at the endoplasmic reticulum (ER). This ER localization was confirmed through co-infiltration with the ER marker SP-mTagBFP2-HDEL (J. Dragdwidge, pers. communication) and subsequent co-localization analysis (**Figure 2A**).

Second, SlFWL7 and -8 were found to localize to both the cytosol and the nucleus. The use of co-infiltrated mScarlet-l marker, a monomeric constitutive red fluorescent protein (Bindel *et al*., 2017), confirmed this localization (**Figure 2B**).

Third, SlFWL2, SlFWL4 and SlFWL5 were targeted to the plasma membrane (PM), as witnessed by staining with FM4.64, a membrane-specific dye (Bolte *et al*., 2004), following plasmolysis using a 0.4 M sorbitol treatment (**Figure 2C**).

We next investigated the topology of these three SlFWLs at the PM using currently available prediction tools. First, the 3D structure of these PM-localized SlFWLs was predicted using ColabFold software (Mirdita *et al*., 2021) based on AlphaFold 2.0 (Jumper *et al*., 2021) and MMseqs2 (Steinegger & Soding, 2017). Second, the insertion in the plasma membrane was modelized using the PPM 3.0 Web Server (Lomize *et al*., 2022) (**Supplementary Fig. S2**). These predictions revealed that the 3D structure of SlFWL2, SlFWL4 and SlFWL5 present similarities to that from FW2.2 (Beauchet *et al*., 2024), such as the absence of any transmembrane domain and their N- and C-terminal parts predicted to be folded on the same side of the protein. These predictions suggest that they do not cross the PM, but are most probably anchored in the PM.

### *In planta* subcellular localization of SlFWL2, SlFWL4 and SlFWL5

Owing to their localization at the PM, we performed an *in planta* analysis of the subcellular localization of SlFWL2, SlFWL4 and SlFWL5, aimed at investigating whether these three SlFWLs are also enriched at PD in tomato, alike FW2.2 (Beauchet *et al*., 2024). For this purpose, we generated stable transgenic lines expressing SlFWL2, SlFWL4 and SlFWL5 fused to YFP at the C-terminal end, under the control of the 35S promoter in the tomato cultivar ‘AC’ (lines referred to as *35S::SlFWL2-YFP*, *35S::SlFWL4-YFP* and *35S::SlFWL5-YFP* plants).

The localization of SlFWL2-YFP, SlFWL4-YFP and SlFWL5-YFP at the PM of tomato leaf cells was confirmed (**Figure 3A**). In addition, SlFWL5-YFP was found to localize at the PM according to a pattern of punctate spots at the cell periphery, suggesting that SlFWL5-YFP was enriched at nanodomains. Staining with aniline blue (AB) to reveal callose deposition as a marker of PD, revealed that only SlFWL5-YFP co-localized with AB, as shown by the overlapping signal intensity plots (**Figure 3A**), thus indicating a localization at PD. Next, we determined the plasmodesmata enrichment ratio, named ‘PD index’, which corresponds to the SlFWLs-YFP fluorescence intensity at PD *vs* that at the cell periphery, to quantify the enrichment of SlFWLs-YFP at PD, as previously described (Beauchet *et al*., 2024) (**Figure 3B**). While the PD index for SlFWL2-YFP and SlFWL4-YFP was equal to 1, a high PD index from 1.5 to 1.7 was measured in leaf cells of *35S::SlFWL5-YFP* plants, thus demonstrating that only SlFWL5 out of the three SlFWLs tested was enriched at PD.

**Fig. 3.**
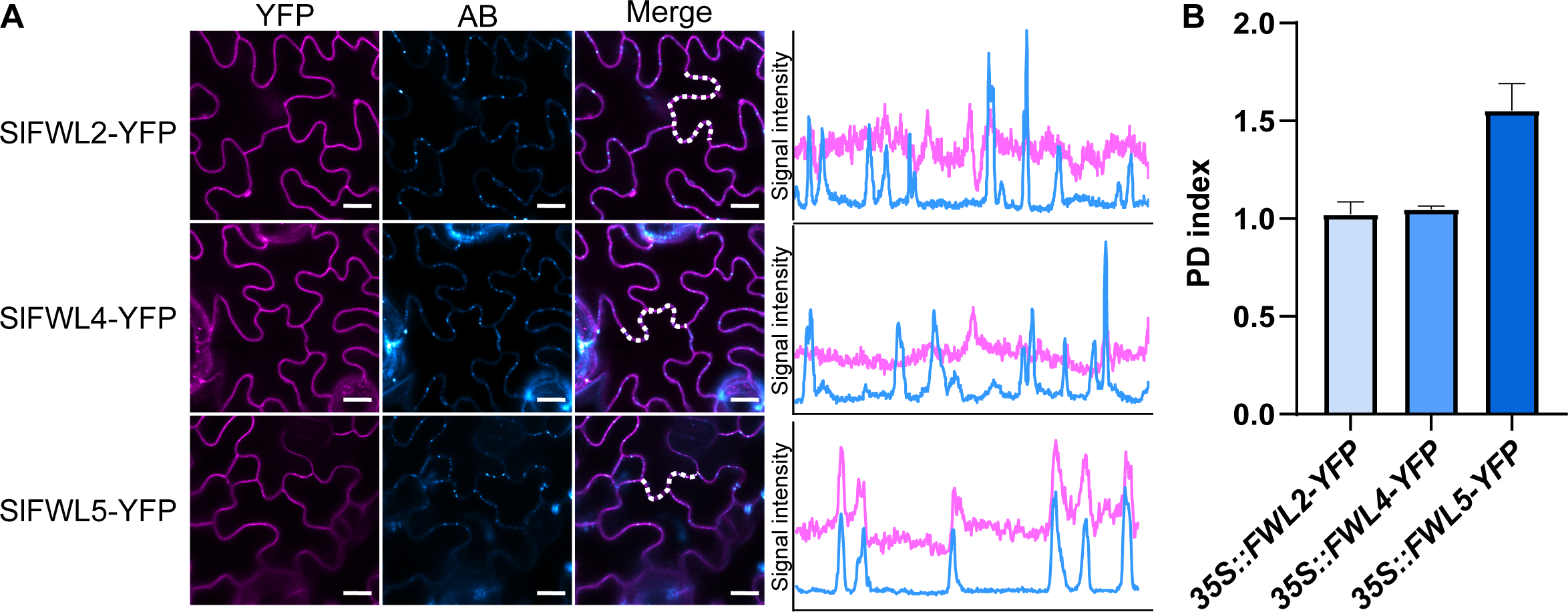
SlFWL5 is enriched at PD. (A) Confocal microscope observations of SlFWL2, SlFWL4 and SlFWL5 localization in tomato leaf cells from *35S::SlFWL2-YFP*, *35S::SlFWL4-YFP*, *35S::SlFWL5-YFP* plants. Scale bar = 10 μm. Intensity plots delineated by the dashed lines are shown for each co-localization pattern. AB, aniline blue. (B) PD index for SlFWL2, SlFWL4 and SlFWL5 localization in leaf cells from *35S::SlFWL2-YF*P, *35S::SlFWL4-YFP and 35S::SlFWL5-YFP* plants. n>20 ROIs from 5 images.

### *In planta* functional analysis of *SlFWL2*, *SlFWL4* and *SlFWL5*

To analyze the function of *SlFWL2*, *SlFWL4* and *SlFWL5* in vegetative and reproductive development, gain-of-function and loss-of-function plants were generated in the tomato cultivar ‘AC’. *SlFWL2*, *SlFWL4* and *SlFWL5* were overexpressed constitutively and ectopically, using the 35S promoter (gain-of-function plants referred to as *35S::SlFWL2, -4* and *-5* respectively). For each gene, three T2 lines were selected with medium- to very high levels of *SlFWL2, -4* and *-5* overexpression in leaves, ranging from 3-fold more to 150-fold more (**Supplementary Fig. S3**). Alternatively, *SlFWL2*, *SlFWL4* and *SlFWL5* were knocked out using the CRISPR/Cas9 technology. Two single-guide RNAs (sgRNAs) per gene were designed close to the start codon, in order to create a frameshift or an early stop codon that would result in a dysfunctional protein (**Supplementary Fig. S4**). For each gene, we selected three independent T0 transgenic lines displaying homozygous mutations, either deletions or insertions, for subsequent phenotyping of vegetative and reproductive development. These loss-of-function plants were referred to as *CR-Slfwl2*, *-4* and *-5* hereafter.

The phenotypic analysis of the different transgenic plants is shown in **Figure 4**. The overexpression of *SlFWL2*, *SlFWL4* and *SlFWL5* did not induce any significant effects on both reproductive and vegetative development. Only one line out of three for the *35S::SlFWL2* and *35S::SlFWL4* constructs, namely *35S::SlFWL2#2* and *35S::SlFWL4#2*, produced smaller fruits (−20% when compared to WT) (**Figure 4A**). No significant impacts could be detected either on root length (**Figure 4B**) or leaf surface (**Figure 4D**), and the sole *35S::SlFWL2#2* line was negatively affected for hypocotyl length, with a reduction in average of 20% when compared to WT (**Figure 4C**).

**Fig. 4.**
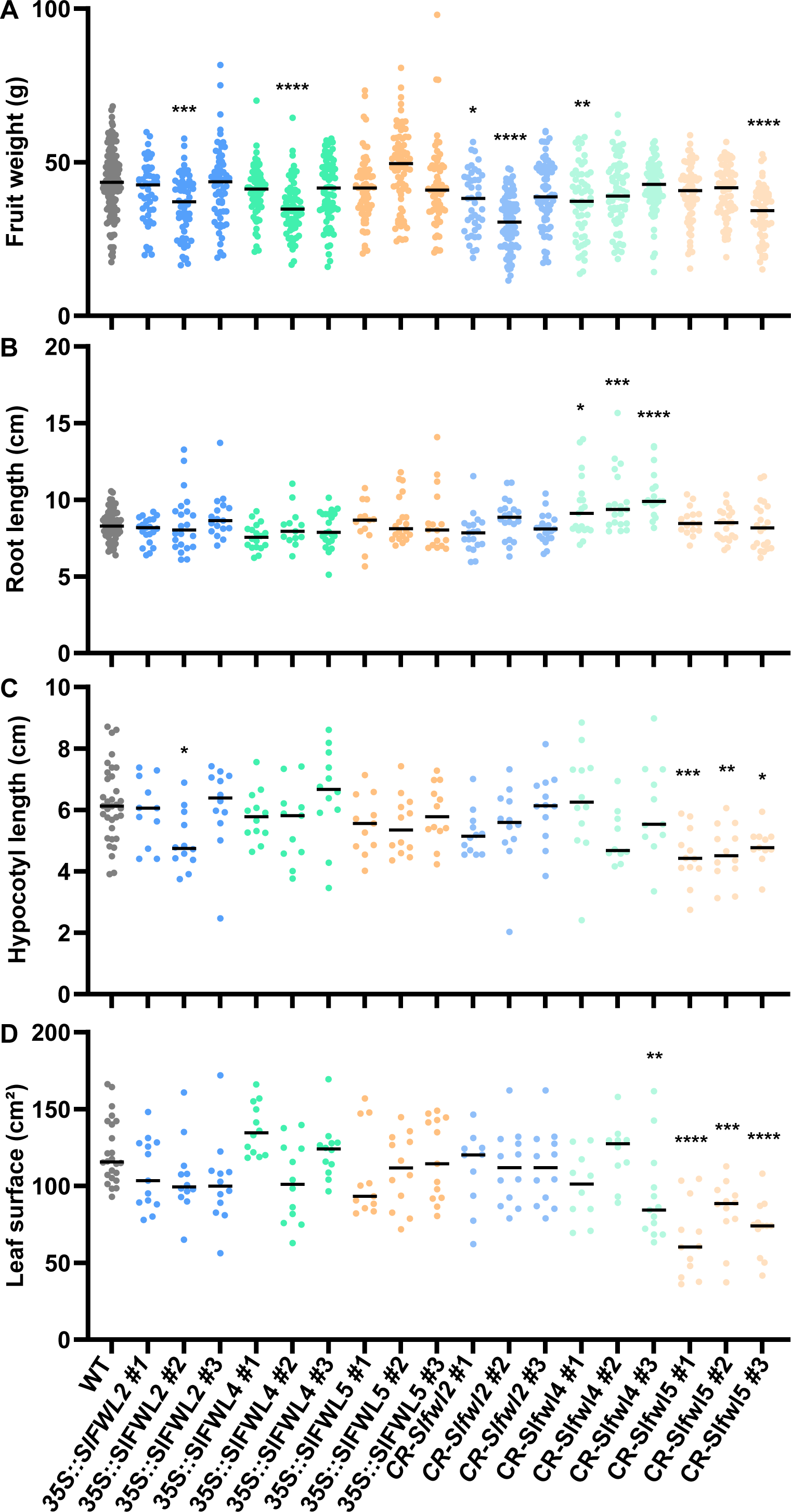
Phenotypic analysis of SlFWL2, SlFWL4 and SlFWL5 gain-of-function (referred to as *35S::SlFWL2*, *35S::SlFWL4* and *35S::SlFWL5*) and loss-of-function plants (referred to as *CR-Slfwl2*, *CR-Slfwl4* and *CR-Slfwl5* compared to WT plants. (A) fruit weight (at red ripe stage); n > 40 fruits from 4 plants per line. (B) Root length (at 17 DAG); n > 10 roots per line. (C) Hypocotyl length (at 17 DAG); n > 10 hypocotyls per line. (D) Mature leaf surface; n > 10 leaves from 4 plants per line. Statistical analysis: Kruskal–Wallis test with post hoc Dunn multiple comparison test. *P < 0.05; ***P < 0.01; ***P < 0.001; ****P < 0.0001.

The most noticeable effects were obtained for CRISPR knockout mutants. Two out of three *SlFWL2* mutant lines (namely *CR-Slfwl2#1* and *CR-fwl2#2*), and the sole *CR-fwl5#3* line displayed a statistically significant reduction in fruit weight (respectively −17%, −42% and −24% in average when compared to WT) (**Figure 4A**). All three *CR-Slfwl4* knockouts lines were affected for root length with an increase from 15% to 23% in average, when compared to WT plants, while *CR-Slfwl2 and -5* lines were not affected (**Figure 4B** and **Supplemental Figure S5**). When the expression of *SlFWL5* is suppressed, the length of the hypocotyl in all three *CR-Slfwl5* lines was reduced from 22% to 38% in average, when compared to WT (**Figure 4C**). More remarkably, all three *CR-Slfwl5* lines displayed a marked reduction in leaf surface (from 32% to 47% in average) (**Figure 4D**), which suggests that the suppression of *SlFWL5* alters the overall vegetative development of aerial organs, as root development was not affected.

To the exception of SlFWL2, these results suggested that vegetative growth and development in tomato are regulated by SlFWL4 in roots and more remarkably, by SlFWL5 in leaves.

### SlFWL5 participates in the control of leaf growth via cell expansion

We then proceeded to a fine phenotyping of the leaf morphology in all three *CR-Slfwl5* lines since the overall leaf surface was reduced compared to the WT. The fully expanded compound leaf of the cultivated tomato cultivar ‘AC’ is made up of a terminal leaflet (TL) and several lateral leaflets attached to the rachis (**Figure 5A**). According to the sequential order and position in the rachis, the lateral leaflets can be further categorized into primary (PL), secondary (SL) and intercalary leaflets (IL).

**Fig. 5.**
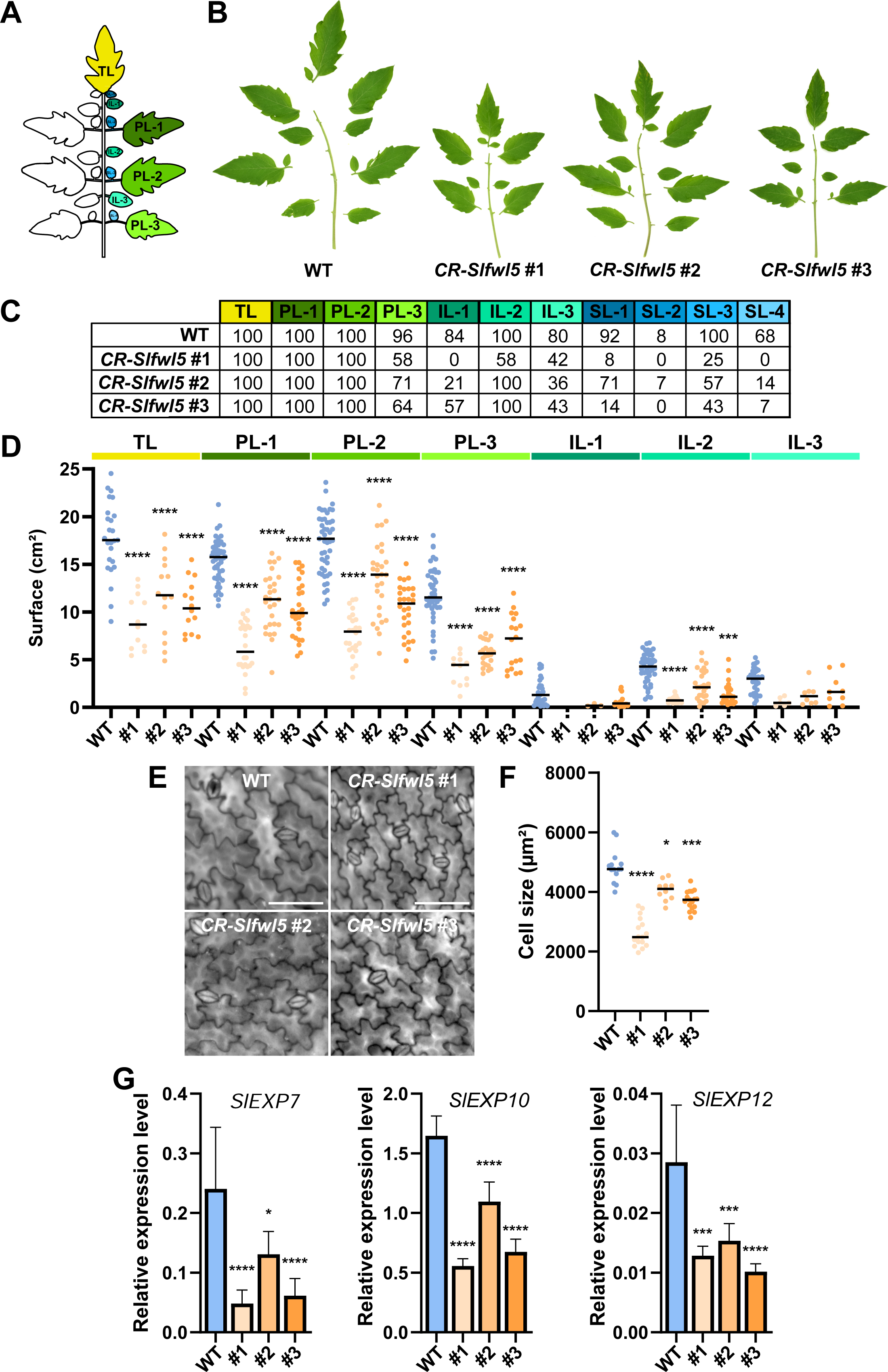
Fine phenotypic analysis of mature leaves from *CR-Slfwl5* plants compared to that of WT. (A) Schematic representation of a mature leaf from wild-type ‘AC’ background defining the terminal leaflet (TL), the primary (PL), secondary (SL) and intercalary leaflets (IL) in their sequential order. (B) Silhouettes of the sixth leaf of 8-week-old CR-Slfwl5 plants. (C) Analysis of leaf composition in *CR-Slfwl5* plants, numbers indicate the percentage of leaves presenting the corresponding leaflet. Color code refers to (A). (D) Determination of the mean TL, PL and IL surface in WT and *CR-Slfwl5* plants. n > 10 leaves from 4 plants per genotype (each dot represents 1 leaflet surface measurement). Statistical analysis: Kruskal–Wallis test with post hoc Dunn multiple comparison test. ***P < 0.001; ****P < 0.0001. (E) Leaf epidermal cells of WT and the three *CR-Slfwl5* plants. Bars = 100 µm. (F) Cell size determination in the leaf epidermis of WT and the three *CR-Slfwl5* plants. Statistical analysis: Kruskal–Wallis test with post hoc Dunn multiple comparison test. *P <0.05; ***P <0.001; *P <0.05; ****P <0.0001. n>12 images measurement from leaves 5 and 6 of 5 independent plants for each line. (G) Expression of SlEXP7, SlEXP10 and SlEXP12 in leaves 3-4 of six-week old WT and *CR-Slfwl5* plants.

When compared to WT leaves, the leaf morphology of all three *CR-Slfwl5* lines was ffected, as the production of PL-3, IL-1, IL-3 and SLs was largely reduced (**Figure 5B**). The *CR-Slfwl5#1* line displayed systematically the most altered leaf composition with the lowest number of leaflets. Not only this reduction in the leaflet number was responsible for the overall reduction in leaf surface observed in all three *CR-Slfwl5* lines compared to WT (**Figure 5C**), but also the reduction in the surface of TL, PL and IL contributed to the phenotype (**Figure 5D**). This reduction in leaflet surface and overall leaf surface in the three *CR-Slfwl5* lines was due to a reduction in cell size ranging from 17 to 46% in average (**Figure 5E** and **5F**). The expression levels of three *EXPANSIN* (*EXP*) genes used as cell expansion markers, namely *SlEXP7*, *-10* and *-12* which are expressed in tomato leaves (Lu *et al*., 2016), were further examined in leaves 3 of six-week old plants of *CR-Slfwl5* lines by RT-qPCR (**Figure 5G**). The expression levels of these three genes were significantly lower in the three mutant lines compared with WT, thus accounting for the alteration of the cell expansion process when the expression of *SlFWL5* is suppressed.

### SlFWL5 pull-down reveals plasma membrane- and plasmodesmata-related proteins

The elucidation of SlFWL5 functional role in leaves and at PD relied on a deeper characterization at the biochemical level. For this purpose, we performed an immunoprecipitation followed by tandem-mass spectrometry (IP-MS/MS) on *35S::SlFWL5-YFP* leaves to identify interacting protein partners of SlFWL5. The relevance of this proteomics approach required to determine the most appropriate leaf developmental stage to harvest, so that the natural interacting protein partners are present in the protein extracts. Hence, the expression level of *SlFWL5* was monitored in the different leaves of 4-week old plants (**Supplemental Figure 6)**. Four-week old plants produced up to 8 leaves: from the newly formed ones, *e.g*. leaves 8 and 7 emerging from the meristem, to the oldest fully-developed ones, *e.g.* leaves 2 and 1. The highest expression of *SlFWL5* was found in leaves 6, 5 and 4, when leaf growth is highly sustained by cell expansion. Protein extracts were then prepared from these three different stages and pooled prior immunoprecipitation.

The IP-MS/MS experiment resulted in the identification of 62 proteins that co-immunoprecipitated with SlFWL5, which were significantly enriched in the *35S::SlFWL5-YFP* sample when compared to *35S::NLS-GFP-GUS* used as a control (**Figure 6A**, **Supplementary Table S3**). A Gene Ontology (GO) term enrichment analysis for the 62 identified proteins was performed (**Figure 6B**), showing that they fall into the three types of GO domains: cellular components (CC), molecular functions (MF) and biological processes (BF). Notably, the CC terms ‘plasma membrane’ and ‘plasmodesmata’ are highly represented. Interestingly, 19 proteins out the 62 identified candidate proteins were found to belong to the FW2.2 co-immunoprecipitation proteome (Beauchet *et al*., 2024), and three of them crossed also with the refined PD proteome from Arabidopsis established by Brault *et al*. (2019), among which was Callose Synthase 10a (SlCalS10a; Solyc03g111570) (**Supplementary Table S4**). It is also noteworthy that among the identified candidate proteins, was a subunit of the Cellulose Synthase complex (SlCesA1; Solyc08g061100), which may relate to cell wall synthesis and cell expansion as regulated by SlFWL5.

**Fig. 6.**
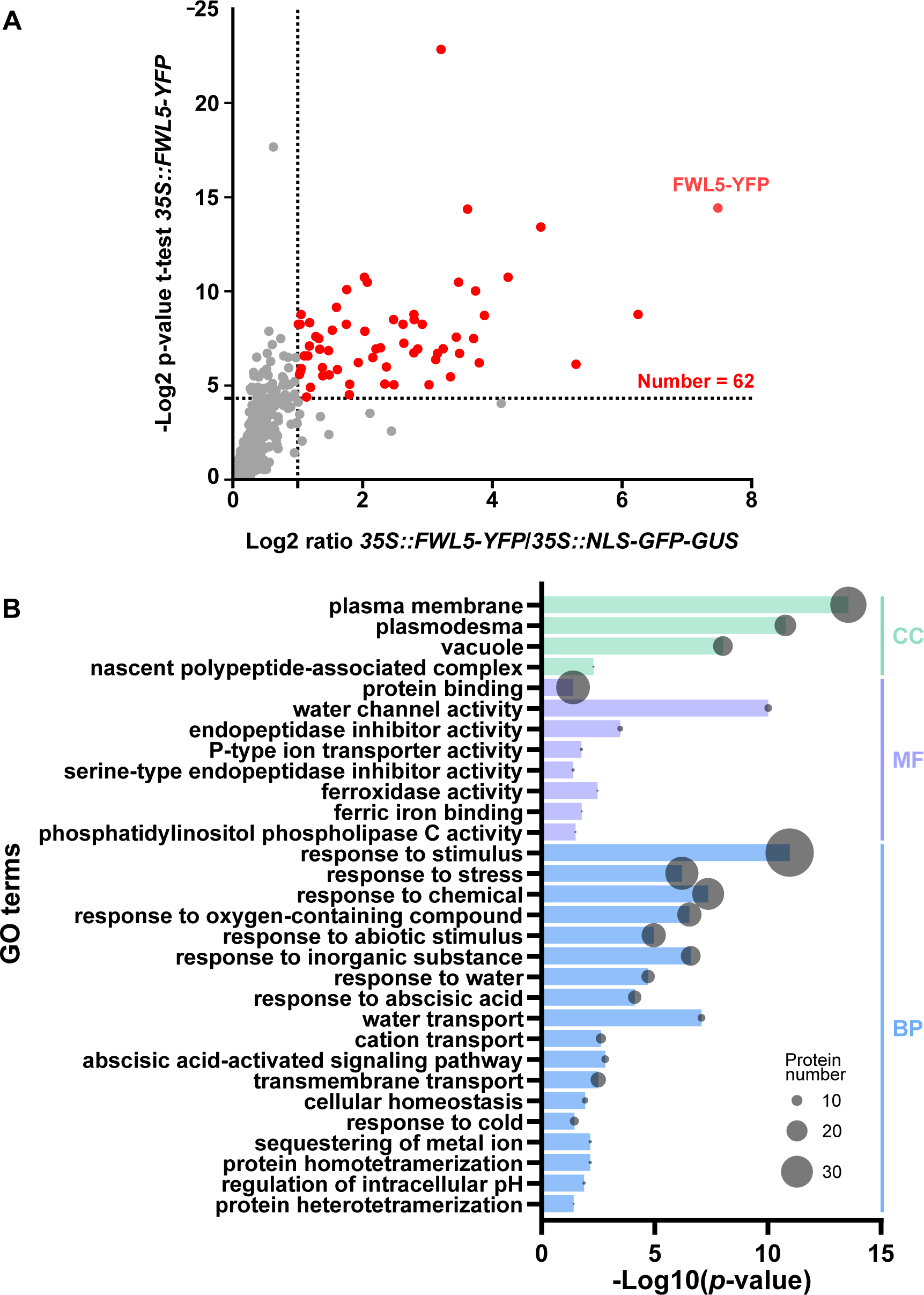
SlFWL5 co-immunoprecipitates with several plasma-membrane and PD-localized proteins. (A) Dot plots showing enriched proteins in *35S::SlFWL5-YFP* IP-MS/MS experiments in young leaves. Red dot indicates significantly enriched protein (based on a Student’s t-test with Benjamini-Hochberg correction P < 0.05 and an enrichment ratio > 2). (B) Gene ontology (GO) enrichment analysis of the identified proteins following SlFWL5 co-immunoprecipitation.

## Discussion

PLAC8 domain-containing proteins in plants belong to the CELL NUMBER REGULATOR/FW2.2-LIKE (CNR/FWL) protein family. Our understanding of the function of CNR/FWL proteins still remains poorly developed, but three distinctive functions have been identified so far (Beauchet *et al*., 2021). These functions relate to (i) calcium uptake and signaling, (ii) metal ion transport and homeostasis, and (iii) organ size determination via the regulation of cell number. These three functions are not exclusive, as they may directly or indirectly influence each other, and examples of mutual actions on plant growth and metal ion transport (Song *et a*l., 2015; Xiong *et al*., 2018; Li *et al*., 2019; Gao *et al*., 2020) or plant growth and calcium signaling (Rosa *et al*., 2017) have been reported. It is noteworthy that a function in improving resistance to a plant pathogen, namely *Xanthomonas oryzae*, has been described alongside to metal ion transport (Li *et al*., 2019).

So far, functional studies of *CNR/FWL* genes are still scarce in the literature. The first reports were dedicated to the functional characterization of *FW2.2* orthologs in important cereal crops such as maize and rice. In maize, the overexpression of *ZmCNR1* resulted in a reduction in overall plant size and organ size (namely tassel, ear and leaf size), thus affecting the plant biomass (Guo *et al*., 2010). In rice, T-DNA insertion mutants of *OsFWL3* and *OsFWL5* displayed an increase in the grain weight and/or plant height (Xu *et al*., 2013; Song *et al*., 2015). Similarly, OsFWL1/OsCNR1 (Ruan *et al*., 2020) and OsFWL4 (Gao *et al*., 2020) were shown to act as negative regulators of rice grain width and weight, and tiller number and plant yield, respectively. In addition, the RNAi silencing of *GmFWL1* expression and the deletion of the conserved PLAC8 domain of *GmFWL3* using CRISPR-Cas9 in soybean resulted in a significant reduction in nodule number in response to rhizobial infection, pointing out at a critical role for GmFWL1 and GmFWL3 in nodule organogenesis (Libault *et al*., 2010; Cervantes-Pérez *et al*., 2024). In all cases, these studies confirmed a role of certain *CNR/FWL* in regulating negatively cell number, thus influencing the organ size and/or development.

Due to the scarcity of data, we have undertaken in the present report a functional analysis of some CNR/FWLs in tomato, as to enrich our knowledge and to decipher their putative role in plant development.

### The CNR/FWL protein family presents a high diversity in protein characteristics, expression pattern and subcellular localization in tomato

The *CNR/FWL* gene family in tomato comprises twenty-one members, showing a large diversity in protein length, ranging from 98 aa (SlFWL1) to 505 aa (SlFWL19). CNR/FWL proteins share as a common feature, the presence of the PLAC8 domain made of two conserved motifs: the more or less divergent Cys-rich motif 1 of the type CLXXXXCPC or CCXXXXCPC, and the more conserved QXXRELK motif 2 (**Figure 1**). Since the CLXXXXCPC motif is present in FW2.2, ZmCNR1, OsFWL1/OsCNR1 and PfCNR1, it would be tempting to address this motif as a signature for CNR/FWL proteins involved in the regulation of organ growth. However, OsFWL4 and OsFWL5 which regulate rice grain weight, harbor the CCXXXXCPC motif. This latter motif has been described as important for conferring the Cd resistance function (Song *et al*., 2004). Interestingly, OsFWL4 and OsFWL5 both enhanced cadmium resistance too in yeast cells, and OsFWL4 acts as a transporter ensuring the translocation of Cd from roots to shoots in rice (Song *et al*., 2015; Xiong *et al*., 2018). Therefore, the sole nature of the PLAC8 Cys-rich motif does not indicate any function with certainty, and even less with the occurrence of different divergent motifs present in the tomato CNR/FWL family members. This was particularly obvious for the three SlFWL proteins we characterized functionally in deeper details, namely SlFWL2, SlFWL4 and SlFWL5, which harbor a CCXXXXCPC, ALXXXXFPC and AVXXXXLPC motif, respectively (**Figure 1**). Indeed, whether SlFWL2, SlFWL4 and SlFWL5 are overexpressed or knocked-out, no significant phenotype could be observed in reproductive development (fruit) (**Figure 4A**). During vegetative development, the sole inactivation of SlFWL4 (*CR-Slfwl4* plants) and SlFWL5 (*CR-Slfwl5* plants) generated marked phenotypes: CR-Slfwl4 lines produced longer roots (**Figure 4B**) and CR-Slfwl5 lines showed reduced hypocotyl length (**Figure 4C**), and a spectacular reduction in leaf size (**Figure 4D** and **5B**). Apart from SlFWL4 which seemed to be preferentially expressed in roots, these phenotypes could not be related with any specific gene expression: SlFWL2 was found more expressed in cotyledons, and SlFWL5 was expressed in all organs, without any preferential expression in leaves (**Supplemental figure S1**).

In the present study, we investigated the subcellular localization of eleven out of twenty SlFWLs (**Figure 2**). A diversity in subcellular localization was observed, as SlFWLs are located according to three distinct patterns: in both the cytosol and the nucleus for SlFWL7 and SlFWL8, at the ER for SlFWL1, -6, -9; -10, -11 and -15, and at the PM for SlFWL2, -4 and -5. Interestingly, a punctate localization in the plasma membrane was observed for SlFWL2 and -4, which may be consistent with a membrane microdomain localization, as demonstrated for GmFWL1 and GmFWL3 in soybean nodules (Qiao et al., 2017; Cervantes-Pérez *et al*., 2024). Therefore, this diversity in subcellular localization might probably be indicative of a diversity in protein function.

To the exception of SlFWL7 and SlFWL8, all the tested SlFWLs were thus found targeted to membranous compartments, most likely in connection with the presence of the hydrophobic PLAC8 domain or TM domains in the case of SlFWL15. From the original structural analysis of AtPCR1 (Song *et al*., 2004), it has long been accepted, and widely and systematically reported in the literature, that the PLAC8 domain in plant CNR/FWLs is composed of two hydrophobic segments (including the CLXXXXCPC or CCXXXXCPC motif), predicted to form transmembrane domains (for a review, see Beauchet *et al*., 2021). However, the use of currently available tools for transmembrane topology prediction revealed that SlFWL2, SlFWL4 and SlFWL5 may not cross the PM in the absence of TM domains, but are likely to be anchored in the outer leaflet of the PM via their PLAC8 hydrophobic domain (**Supplemental Figure S2**), Such a topology would thus be very similar to that demonstrated experimentally for FW2.2 (Beauchet *et al*., 2024).

### SlFWL5 regulates leaf size in tomato via cell expansion

Unexpectedly, we could clearly demonstrate that the leaf growth phenotype in *CR-Slfwl5* lines does not originate from an impairment in cell division but from an impairment in cell expansion, as cell size in all three lines was greatly reduced and leaf-expressed *EXPANSIN* genes were significantly repressed (**Figure 5E-G**). The contribution of *EXPANSIN* gene expression in modulating leaf growth is a long-standing observation (Cho and Cosgrove, 2000; Goh *et al*., 2012), and is part of the complex molecular and hormonal network regulating leaf size (Gonzalez *et al*., 2012).

Not only the inactivation of *SlFWL5* affected leaf growth, but it altered also leaf morphology into simpler leaves, displaying a reduced number of the third primary leaflet, of secondary leaflets and intercalary leaflets (**Figure 5C-D**). This phenotype is fully relevant with hormonal control of compound leaf morphogenesis and differentiation in tomato (Bar and Ori, 2015). Indeed, gibberellin (GA) negatively regulates leaf complexity in tomato, as increased GA levels or GA signaling accelerate leaf maturation and induce simpler leaves than in wild type (Jasinski *et al*., 2008; Yanai *et al*., 2011). In addition, GA and cytokinin (CK) present a mutual antagonistic interaction during tomato leaf development (Fleishon *et al*., 2011). How to relate the molecular origin of the leaf phenotype in *CR-Slfwl5* plants and phytohormone signaling remains to be established, but the following step in the functional characterization of SlFWL5 may provide new lines of research.

### SlFWL5 is enriched at plasmodesmata, within a protein complex composed of plasmodesmata- and plasma membrane-specific proteins

We have shown that SlFWL5 is enriched at PD (**Figure 3**), where it probably participates in a protein complex composed of plasma membrane- and plasmodesmata-specific proteins (**Figure 6**). Interestingly, a significant number of candidate proteins that co-immunoprecipitated with SlFWL5 were found enriched at PD. This finding suggests that SlFWL5 may have a function in cell-to-cell communication mechanisms for the control of leaf size and morphology. SlFWL5 is able to interact with Callose Synthase 10a (CalS10a; Solyc03g111370) which contributes to callose homeostasis at the cell plate and at PD, thereby regulating cytokinesis and the symplastic molecular exchanges between neighboring cells via the permeability of PD (Saatian *et al*. 2018). CalS10 is required for normal plant development, as silencing *CalS10* results in retarded growth: plants display dwarfism, with smaller stem and smaller leaves (Töller *et al*. 2008), a phenotype originating from either impaired cell division or cell expansion. *CR-Slfwl5* loss-of-function plants displayed a similar phenotype, *i.e.* a reduced hypocotyl length and leaf surface, due to impaired cell expansion. Interestingly, SlFWL5 interacts also with SlCesA1, a subunit of the Cellulose Synthase complex. Cellulose is the most abundant β-glucan polysaccharide of the plant cell wall, which self-assemble as microfibrils to strengthen the cell wall and contribute to the direction of cell growth during the cell expansion process (Schneider *et al*., 2016). Both cellulose and callose are synthesized at the PM by the large respective synthase complexes, cellulose synthases and callose synthases, and the formation of cellulose/callose networks has been demonstrated, especially as a layered cellulose/callose architecture in cell walls of epidermal leaf cells (Falter *et al*., 2015). Whether the SlFWL5 function in controlling leaf growth requires a regulatory role on Callose- and Cellulose synthases during cell expansion remains an exciting matter of future investigation.

As key elements of the cell-to-cell communication machinery, PDs are implicated in processes guaranteeing the collaborative function of the cells, developmental and patterning events (Petit *et al*., 2020), enabling the intercellular trafficking of different mobile signaling molecules, such as small RNAs, metabolites, hormones and transcription factors (TFs) (Wu and Gallagher, 2012; Brunkard et al., 2013). Being localized at PD, SlFWL5 may harbor a regulatory role on plasmodesmata activity. Hence, we can hypothesize that SlFWL5 function is to regulate the cell-to-cell movement of molecules/signals promoting leaf cell expansion and leaflet initiation. Obviously, the identification of such signaling molecules belonging to transcriptional networks involved in pathways affecting cell growth (Davière and Achard, 2013), and leaf growth (Gonzalez *et al*., 2012; Bar and Ori, 2014), would represent a giant leap in understanding the SlFWL5 functional role at PD.

In conclusion, we are far from deciphering the functional complexity of the CNR/FWL protein family during plant and organ development. Unexpectedly, we have been able to demonstrate in this report that SlFWL5 contributes to the regulation of cell expansion during tomato leaf development. Therefore, this study may lead us to revise our perception of the CNR/FWL function in regulating organ size. Whatever the mechanism is involved, cell division or cell expansion control, what may really matter is the potential function in cell-to-cell communication molecules as to regulate organ growth.

## Supplementary Data

The following supplementary data are available at JXB online.

**Fig. S1.** SlFWLs expression analysis in different organs of tomato plants and during fruit development.

**Fig. S2.** Structure and membrane insertion predictions of SlFWL2, SlFWL4 and SlFWL5 compared to that of FW2.2.

**Fig. S3.** RT-qPCR analysis of SlFWL2, SlFWL4 and SlFWL5 expression in leaves of their respective overexpressing lines.

**Fig. S4.** CRISPR/Cas9-induced mutations producing truncated versions of SlFWL2, SlFWL4 and SlFWL5.

**Fig. S5.** Illustration of root phenotype in WT and *CR-Slfwl4* loss-of-function tomato plants.

**Fig. S6.** SlFWL5 expression in the course of leaf development in six weeks-old WT plants.

**Table S1.** List of primers used for constructs and RT-qPCR analysis.

**Table S2.** List of Gateway vectors used for constructs.

**Table S3.** Raw datasets from IP-MS/MS following SlFWL5-YFP co-immunoprecipitation.

**Table S4.** List of candidate proteins identified as interactors of SlFWL5.

## Acknowledgements

We express our deepest thanks to Isabelle Atienza, Aurélie Honoré and Valérie Rouyère, for taking care of the plant culture in the greenhouse. We thank Jonathan Dragdwidge (VIB-UGent for Plant Systems Biology, Ghent University, Belgium) for sharing the unpublished *pH3.3::SP-mTagBFP2-HDEL* and *pUBI10::mScarlet-I* plasmids. Mass spectrometry experiments were carried out using the facilities of the Montpellier Proteomics Platform (PPM, BioCampus Montpellier, France). The microscopy analyses were performed in the Bordeaux Imaging Center, a service unit of the CNRS-INSERM and Bordeaux University, member of the national infrastructure France BioImaging, and supported by the French National Research Agency (ANR-10-INBS-04), with the great help of Lysiane Brocard.

## Author contributions

N.G., C.C. and N.B. conceived the project and designed the research. A.B., L.E., L.B. and N.B. performed the research. V.R. performed the IP-MS-MS proteomics experiments. All authors analyzed and discussed the results. A.B., N.G., C.C. and N.B. wrote the manuscript with input from the other authors.

## Conflict of interest

The authors declare that they have no conflicts of interest.

## Funding

This work was carried out with the financial support of the French Agence Nationale de la Recherche (grant no. ANR-20-CE20-0002), the GPR Bordeaux Plant Sciences in the framework of the IdEX Bordeaux University ‘Investments for the Future’ program, and the French Ministère de l’Enseignement Supérieur et de la Recherche (PhD grant to A. Beauchet).

## Data availability

The Mass Spectrometry Proteomics data underlying this article have been deposited to the ProteomeXchange Consortium via the PRIDE (Perez-Riverol *et al*., 2022) partner repository with the dataset identifier XXXX.

## Notes

### Competing Interest Statement

The authors have declared no competing interest.

## References

Alpert KB, Grandillo S, Tanksley SD. 1995. fw 2.2:a major QTL controlling fruit weight is common to both red- and green-fruited tomato species. Theoretical and Applied Genetics 91, 994–1000.

Beauchet A, Gévaudant F, Gonzalez N, Chevalier C. 2021. In search of the still unknown function of FW2.2/CELL NUMBER REGULATOR, a major regulator of fruit size in tomato. Journal of Experimental Botany 72, 5300–5311.

Beauchet A, Bollier N, Grison M, Rofidal V, Gévaudant F, Bayer E, Gonzalez N, Chevalier C. 2024. The CELL NUMBER REGULATOR FW2.2 protein regulates cell-to-cell communication in tomato by modulating callose deposition at plasmodesmata. Plant Physiology 196, 883–901.

Bindels DS, Haarbosch L, van Weeren L et al. 2017. mScarlet: a bright monomeric red fluorescent protein for cellular imaging. Nature Methods 14, 53–56.

Bolte S, Talbot C, Boutte Y, Catrice O, Read ND, Satiat-Jeunemaitre B. 2004. FM-dyes as experimental probes for dissecting vesicle trafficking in living plant cells. Journal of microscopy. 214, 159–173.

Brault ML, Petit J, Immel F et al. (2019). Multiple C2 domains and transmembrane region proteins (MCTPs) tether membranes at plasmodesmata. EMBO Reports 20, e47182.

Brunkard, JO, Anne M, Runkel AM, Zambryski PC. 2013. Plasmodesmata dynamics are coordinated by intracellular signaling pathways. Current Opinion in Plant Biology 16, 614–620.

Cabreira-Cagliari C, Dias NC, Bohn B, Fagundes DGDS, Margis-Pinheiro M, Bodanese-Zanettini MH, Cagliari A. 2018. Revising the PLAC8 gene family: from a central role in differentiation, proliferation, and apoptosis in mammals to a multifunctional role in plants. Genome 61, 857–865.

Cervantes-Pérez SA, Zogli P, Amini S, et al. 2024. Single-cell transcriptome atlases of soybean root and mature nodule reveal new regulatory programs controlling the nodulation process. Plant Communications 5, 100984.

Cho HT, Cosgrove DJ. 2000. Altered expression of expansin modulates leaf growth and pedicel abscission in *Arabidopsis thaliana*. Proceedings of the National Academy of Sciences, USA 97, 9783–9788.

Cong B, Liu J, Tanksley SD. 2002. Natural alleles at a tomato fruit size quantitative trait locus differ by heterochronic regulatory mutations. Proceedings of the National Academy of Sciences, USA 99, 13606–13611.

Dahan Y, Rosenfeld R, Zadiranov V, Irihimovitch V. 2010. A proposed conserved role for an avocado FW2.2-like gene as a negative regulator of fruit cell division. Planta 232, 663–676.

Davière JM, Achard P. 2013. Gibberellin signaling in plants. Development 140, 1147–1151.

Falter C, Zwikowics C, Eggert D, Blümke A, Naumann M, Wolff K, Ellinger D, Reimer R, Voigt CA. 2015. Glucanocellulosic ethanol: the undiscovered biofuel potential in energy crops and marine biomass. Scientific Reports 5, 13722.

Fleishon S, Shani E, Ori N, Weiss D. 2011. Negative reciprocal interactions between gibberellin and cytokinin in tomato. New Phytologist 190, 609–617.

Frary A, Nesbitt TC, Grandillo S, Knaap E, Cong B, Liu J, Meller J, Elber R, Alpert KB, Tanksley SD. 2000. fw2.2: a quantitative trait locus key to the evolution of tomato fruit size. Science 289, 85–88.

Galaviz-Hernandez C, Stagg C, de Ridder G, Tanaka TS, Ko MSH, Schlessinger D, Nagaraja R. 2003. Plac8 and Plac9, novel placental-enriched genes identified through microarray analysis. Gene 309, 81–89.

Gao Q, Li G, Sun H, Xu M, Wang H, Ji J, Wang D, Yuan C, Zhao X. 2020. Targeted Mutagenesis of the Rice FW 2.2-Like Gene Family Using the CRISPR/Cas9 System Reveals OsFWL4 as a Regulator of Tiller Number and Plant Yield in Rice. International Journal of Molecular Sciences 21, 809.

Gao Q, Liu L, Zhou H et al. 2021. Mutation in OsFWL7 affects Cadmium and micronutrient metal accumulation in rice. International Journal of Molecular Sciences 22, 12583.

Goh HH, Sloan J, Dorca-Fornell C, Fleming A. 2012. Inducible repression of multiple expansin genes leads to growth suppression during leaf development. Plant Physiology 159, 1759–1770.

Gonzalez N, Vanhaeren H, Inzé D. 2012. Leaf size control: complex coordination of cell division and expansion. Trends in Plant Science 17, 332–240.

Grison MS, Kirk P, Brault ML, Wu XN, Schulze WX, Benitez-Alfonso Y, Immel F, Bayer EM. 2019. Plasma membrane-associated receptor-like kinases relocalize to plasmodesmata in response to osmotic stress. Plant Physiol. 181, 142–160.

Guo M, Rupe MA, Dieter JA, Zou J, Spielbauer D, Duncan KE, Howard RJ, Hou Z, Simmons CR. 2010. Cell Number Regulator1 affects plant and organ size in maize: implications for crop yield enhancement and heterosis. Plant Cell 22, 1057–1073.

Hallgren J, Tsirigos KD, Pedersen MD, et al., 2022. DeepTMHMM predicts alpha and beta transmembrane proteins using deep neural networks. BioRxiv, 2022–04.

Jasinski S, Tattersall A, Piazza P, Hay A, Martinez-Garcia JF, Schmitz G, Theres K, McCormick S, Tsiantis M. 2008. *PROCERA* encodes a DELLA protein that mediates control of dissected leaf form in tomato. Plant Journal 56, 603–612.

Jumper J, Evans R, Pritzel A, et al. 2021. Highly accurate protein structure prediction with AlphaFold. Nature, 596, 583–589.

Li Z, He C. 2015. Physalis floridana Cell Number Regulator1 encodes a cell membrane-anchored modulator of cell cycle and negatively controls fruit size. Journal of Experimental Botany 66, 257–270.

Li B, Sun S, Gao X, Wu M, Deng Y, Zhang Q, Li X, Xiao J, Ke Y, Wang S. 2019. Overexpression a “fruit-weight 2.2-like” gene OsFWL5 improves rice resistance. Rice 12, 51.

Libault M, Zhang X-C, Govindarajulu M, Qiu J, Ong YT, Brechenmacher L, Berg RH, Hurley-Sommer A, Taylor CG, Stacey G. 2010. A member of the highly conserved FWL (tomato FW2.2-like) gene family is essential for soybean nodule organogenesis. Plant Journal 62, 852–864.

Lomize AL, Todd SC, Pogozheva ID. 2022. Spatial arrangement of proteins in planar and curved membranes by PPM 3.0. Protein Science 31, 209–220.

Liu J, Cong B, Tanksley SD. 2003. Generation and analysis of an artificial gene dosage series in tomato to study the mechanisms by which the cloned quantitative trait locus fw2.2 controls fruit size. Plant Physiology 132, 292–299.

Lu Y, Liu l, Wang X, HanZ, Bo Ouyang B, Zhang Z, Li H. 2016. Genome-wide identification and expression analysis of the expansin gene family in tomato. Molecular Genetics and Genomics 291, 597–608.

Mao M, Cheng Y, Yang J et al. 2021. Multifaced roles of PLAC8 in cancer. Biomarker Research 9, 73.

Marshall OJ. PerlPrimer: cross-platform, graphical primer design for standard, bisulphite and real-time PCR. 2004. Bioinformatics 20, 2471–2472

Mirdita M, Schütze K, Moriwaki Y, Heo L, Ovchinnikov S, & Steinegger, M. 2022. ColabFold: making protein folding accessible to all. Nature Methods 19, 679–682.

Nesbitt TC, Tanksley SD. 2001. fw2.2 Directly Affects the Size of Developing Tomato Fruit, with Secondary Effects on Fruit Number and Photosynthate Distribution. Plant Physiology 127, 575–583.

Perez-Riverol Y, Bai J, Bandla C, García-Seisdedos D, Hewapathirana S, Kamatchinathan S, Kundu DJ, Prakash A, Frericks-Zipper A, Eisenacher M, et al. 2022. The PRIDE database resources in 2022: a hub for mass spectrometry-based proteomics evidences. Nucleic Acids Research 50, D543–D552.

Petit JD, Li ZP, Nicolas WJ, Grison M, Bayer E. 2020. Dare to change, the dynamics behind plasmodesmata-mediated cell-to-cell communication. Current Opinion in Plant Biology 53, 80–89.

Qiao Z, Brechenmacher L, Smith B, Strout GW, Mangin W, Taylor C, Russell SD, Stacey G, Libault M. 2017. The GmFWL1 (FW2-2-like) nodulation gene encodes a plasma membrane microdomain-associated protein. Plant, Cell & Environment 40, 1442–1455.

Ran C, Zhang Y, Chang F, Yang X, Liu Y, Wang Q, Zhu W. 2023. Genome-wide analyses of *SlFWL* family genes and their expression profiles under cold, heat, salt and drought stress in tomato. International Journal of Molecular Sciences 24, 11783.

Rosa M, Abraham-Juárez MJ, Lewis MW et al. 2017. The maize mid-complementing activity homolog cell number regulator13/narrow odd dwarf coordinates organ growth and tissue patterning. The Plant Cell, 29, 474–490.

Ruan B, Shang L, Zhang B, Hu J, Wang Y, Lin H, Zhang A, Liu C, Peng Y, Zhu L, et al. 2020. Natural variation in the promoter of TGW2 determines grain width and weight in rice. New Phytologist 227, 629–640.

Saatian B, Austin RS, Tian G, Chen C, Nguyen V, Kohalmi SE, Geelen D, Cui Y. 2018. Analysis of a novel mutant allele of *GSL8* reveals its key roles in cytokinesis and symplastic trafficking in Arabidopsis. BMC Plant Biology 18, 295.

Schneider R, Hanak T, Persson S, Voigt CA. 2016. Cellulose and callose synthesis and organization in focus, what’s new? Current Opinion in Plant Biology 34, 9–16.

Song W-Y, Martinoia E, Lee J, Kim D, Kim D-Y, Vogt E, Shim D, Choi KS, Hwang I, Lee Y. 2004. A novel family of cys-rich membrane proteins mediates cadmium resistance in Arabidopsis. Plant Physiology 135, 1027–1039.

Song WY, Choi KS, Kim DY et al. 2010. Arabidopsis PCR2 Is a zinc exporter involved in both zinc extrusion and long-distance zinc transport. Plant Cell 22, 2237–2252.

Song WY, Lee HS, Jin SR, Ko D, Martinoia E, Lee Y, An G, Ahn SN. 2015. Rice PCR1 influences grain weight and Zn accumulation in grains. Plant Cell & Environment 38, 2327–2339.

Steinegger M, & Söding J. 2017. MMseqs2 enables sensitive protein sequence searching for the analysis of massive data sets. Nature biotechnology, 35, 1026–1028.

Swinnen G, Mauxion JP, Baekelandt A et al. 2022. SlKIX8 and SlKIX9 are negative regulators of leaf and fruit growth in tomato. Plant physiology 188, 382–396.

Tamura K, Stecher G, Kumar S. 2021. MEGA11: Molecular Evolutionary Genetics Analysis version 11. Molecular Biology and Evolution 38, 3022–3027.

Thibivilliers S, Farmer A, Libault M. 2020. Biological and Cellular Functions of the Microdomain-Associated FWL/CNR Protein Family in Plants. Plants 9, 377.

Töller A, Brownfield L, Neu C, Twell D, Schulze-Lefer P. 2008. Dual function of Arabidopsis glucan synthase-like genes GSL8 and GSL10 in male gametophyte development and plant growth.

Van Bel M, Silvestri F, Weitz EM et al. 2022. PLAZA 5.0: Extending the scope and power of comparative and functional genomics in plants. Nucleic Acids Research 50, 1468–1474.

Wu S, Gallagher KL. 2012. Transcription factors on the move. Current Opinion in Plant Biology 15, 645–651.

Xiong W, Wang P, Yan T, Cao B, Xu J, Liu D, Luo M. 2018. The rice “fruitlZlweight 2.2lZllike” gene family member OsFWL4 is involved in the translocation of cadmium from roots to shoots. Planta 247, 1247–1260.

Xu J, Xiong W, Cao B, Yan T, Luo T, Fan T, Luo M. 2013. Molecular characterization and functional analysis of ‘fruit-weight 2.2-like’ gene family in rice. Planta 238, 643–655.

Yanai O, Shani E, Russ D, Ori N. 2011. Gibberellin partly mediates LANCEOLATE activity in tomato. Plant Journal 68, 571–582

Zsögön A, Čermák T, Rezende Naves E, et al. 2018 De novo domestication of wild tomato using genome editing. Nature Biotechnology 36, 1211–1216.

